# A novel mechanism of morphogenetic transition in *Schizosaccharomyces pombe* dependent on the MAPK Spc1 and the transcription factor Atf1 associated with sensing of optimal growth environment

**DOI:** 10.1101/2025.05.17.654634

**Authors:** Suchismita Datta, Suparna Dutta, Jayshikha Banerjee, Geetanjali Sundaram

**Author notes:** Corresponding author;, Contact No. 9433191982. These authors have contributed equally to the work (Suparna Dutta, Jayshikha Banerjee). **Conflicts of Interest**: The authors declare no competing or financial interest.

## Abstract

Fission yeast *Schizosaccharomyces pombe* are rod shaped eukaryotic cells that grow by tip elongation and divide medially by formation of cell wall septum. The coordination between cell length and cell cycle phase transitions in this simple eukaryote makes it an excellent model system for studying morphogenetic events associated with both intrinsic and extrinsic disturbances in cell cycle regulation. Cell cycle progression in *S. pombe* is dependent on the coordinated functions of two bZIP family transcription factors Atf1 and Pcr1. In an attempt to understand this coordination, we stumbled upon the startling observation that overexpression of Atf1 in cells lacking Pcr1 led to drastic morphogenetic alteration whereby the cells became hyper elongated. In this report we present the evidences of dependence of this phenotype on metabolic status of the cell. We also show that the phenotype is dependent on the activity of the <u>M</u>itogen <u>A</u>ctivated <u>P</u>rotein <u>K</u>inase, Spc1 and that it is associated with the global gene expression alterations of cells resulting from the perturbed balance of Atf1 and Pcr1 activities. We found that change in the carbon source, alterations in iron concentration and the redox environment of the cell can affect the extent of morphogenesis, to the extent of even abolishing it at times. Our results also indicate that this phenotype has a strong association with maintenance of optimum growth conditions of the cell. Taken together, our observations reveal a completely novel mechanism of morphogenetic transition in fission yeast involving the MAPK Spc1 and its downstream effector molecule Atf1, which can be tuned by the cell’s metabolic environment.

## Introduction

Yeasts are widely used as model organism because they are unicellular with genome amenability which aids in detailed investigation of several cellular processes of a eukaryotic system. Several yeasts species exhibit dimorphism in their life cycle as a part of their response to conditions of their external environment e.g. limited nitrogen reserves can induce pseudohyphal differentiation in budding yeast while [1,2,3] serum is seen to induce yeast to hyphal transition in pathogen species like *Candida albicans* [4].

Dimorphism is also seen in homothallic *S. pombe* cells. Naturally occurring *S. pombe* cells are homothallic (h^90^) and can switch between the two mating types-h^+^ and h^-^ [5] thereby allowing them to initiate sexual reproduction when exposed to adverse conditions. They undergo meiosis and form spores which can then germinate after favorable growth conditions resume. *S. pombe* is known to switch its mating forms in response to nitrogen starvation [3] which is under the regulation of the MAPK pathway [6] and the cAMP responsive pathways [7]. These homothallic *S. pombe* cells sometimes exhibit transition from yeast to filamentous form in specific conditions like increase in ammonium [8] or lack of nitrogen [9] and are regulated by cAMP pathways.

MAP kinase cascades are ancient and conserved pathways found in eukaryotes from yeast to humans [10] and are known to regulate cytoplasmic and nuclear events in response to external stimuli [11,12,13]. In *Schizosaccharomyces pombe* there are three different MAPK cascades studied so far. The Spk1 MAPK pathway required for mating and sporulation [14, 15, 16, 17, 18]. Cell wall integrity, cell morphology and response to ion homeostasis is brought about by the Pmk1/Spm1 pathway, another MAPK cascade [19]. The mechanism of regulation of meiosis and sporulation in *S. pombe* is also dependent on the MAPK Spc1 and its downstream effector Atf1, which is a bZIP domain containing transcription factor [20]. The main role of the Spc1 MAPK pathway is stress response regulation. This three tiered kinase cascade consists of Win1 (MAPKKK), Wis1 (MAPKK) and Spc1 (MAPK) which senses the stress stimuli and regulates expression of specific stress-induced genes in response to oxidative stress, osmotic stress, ion disbalance and nutrient limitation [21, 22, 23, 24]. These effects involve global gene expression changes which are mainly brought about by its downstream effector molecule Atf1 [20, 25, 26]. This bZIP domain containing transcription factor gets targeted to stress responsive promoters via a mechanism that involves its binding partner Pcr1, which is also a bZIP domain containing transcription factor [27, 28, 29]. Atf1 is also important for meiotic recombination, sexual differentiation and heterochromatin formation [27, 30, 31, 32, 33]. It is also necessary for the accumulation of the cells in G1 during nitrogen starvation [20] and also associated with the spindle orientation checkpoint [34]. It also regulates the anaphase-promoting complex [35]. Studies in our lab have established Atf1 to be an important regulator of both mitotic entry [36, 37, 38] and metaphase to anaphase transition [39] in *S. pombe*. We have also shown that the distinction between stress responsive and cell cycle accelerating functions of Atf1 is also mediated by Pcr1 which can selectively inhibit the recruitment of Atf1 on cell cycle related targets [38].

This study was undertaken to understand how the interplay between Atf1 and Pcr1 can affect the life cycle of homothallic *S. pombe* cells. To this end we adopted the strategy of overexpressing Atf1 in h^90^ *S. pombe* cells both in absence and presence of Pcr1. To our surprise we found that Atf1 overexpression in absence of Pcr1 triggered the morphogenetic transition of these yeast cells to the filamentous form even during favourable growth conditions. This observation implied that Atf1 and Pcr1 can be important regulators of dimorphism in homothallic *S. pombe* cells and hence we proceeded towards further characterization of the conditions associated with this morphogenesis. Our results show that the yeast to filamentous form morphogenesis is tightly linked with the nutritional status and metabolic conditions of the cell. We also show that the Spc1 activity, phosphorylation status of Atf1 and gene expression changes is brought about by Atf1 overexpression.

Our results illuminate how alternations in iron homeostasis and changes in carbon source can have an important bearing upon the morphology of the homothallic *S. pombe* cells. We also show that the metabolic tuning of MAPK Spc1 is associated with the dimorphism observed in *S. pombe*.

## Results and Discussion

### Perturbation of Atf1-Pcr1 balance leads to significant morphogenetic changes in *S. pombe*

Earlier studies in our lab have shown that the bZIP transcription factors, Atf1 and Pcr1 have antagonistic influences on cell cycle phase transitions. To further understand this regulatory mechanism we decided to study the cell cycle progression in cells where the relative levels of Atf1 and Pcr1 are changed via genetic modification. Surprisingly we found a drastic morphogenetic transition when we overexpressed Atf1 in *Δpcr1* mutants. These cells showed a significantly elongated filamentous morphology (Figure 1A) in contrast to *wt* cells overexpressing Atf1, where the cells become shorter due to accelerated mitotic entry. The remarkably different outcomes of Atf1 overexpression in *wt* and *Δpcr1* cells were interesting and revealed that these two transcription factors might have a very significant role to play in cell morphology and fate determination. These elongated cells were observed to be mononucleated but the nuclear morphology showed aberrations which indicated the possible presence of mitotic defects in these conditions (Figure 1B). Septum morphology seemed normal although expectedly septa were found to be less frequent in a population of these cells (Figure 1B). The growth rate (Figure 1C) as well as viability (Figure 1D) of these cells was much less compared to *wt* cells overexpressing Atf1 as well as *Δpcr1* cells transformed with the empty vector.

**Figure 1.**
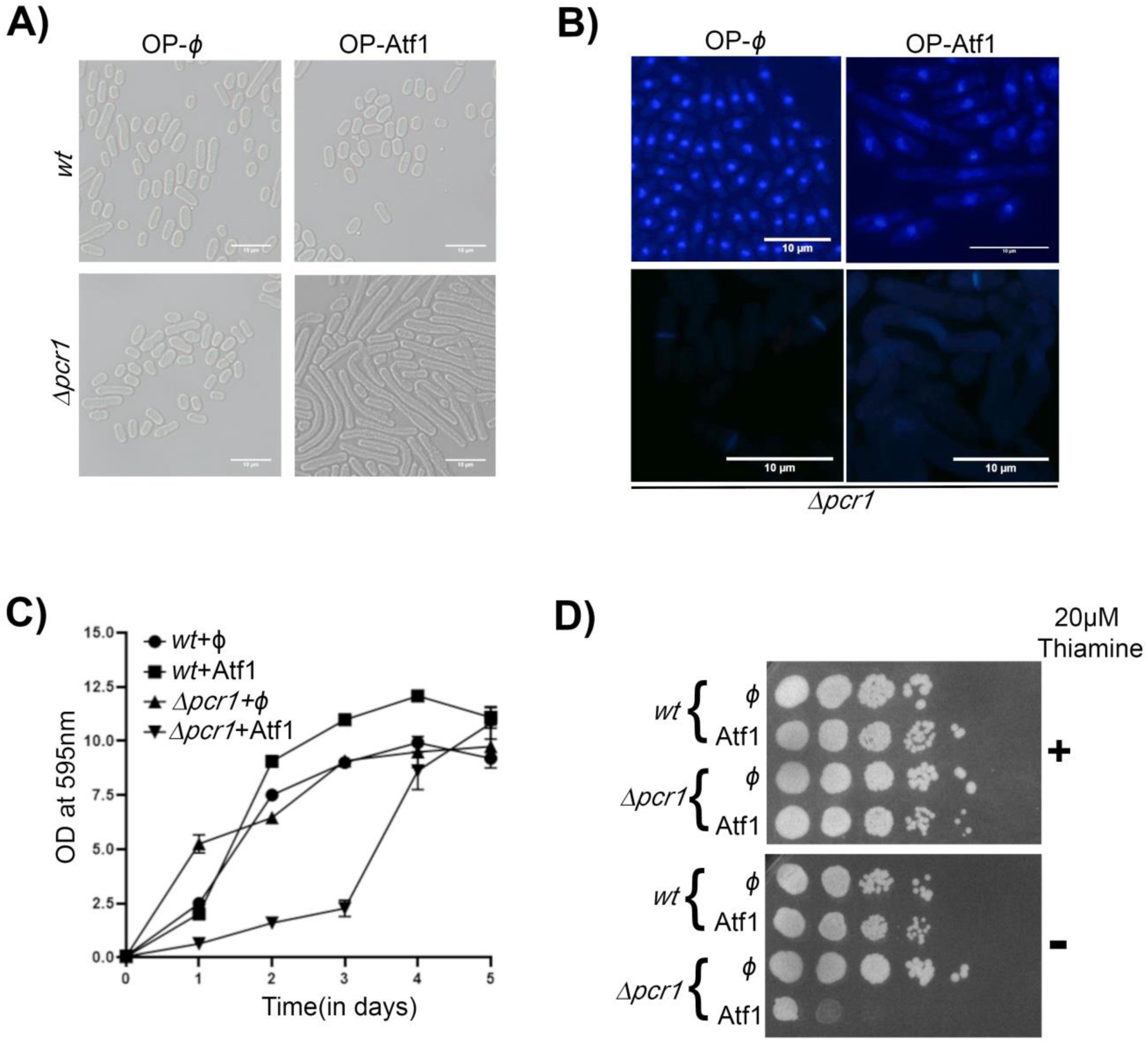
Perturbation of Atf1-Pcr1 balance leads to significant morphogenetic changes in *S. pombe*. (A) Bright-field images of *wt and Δpcr1* cells transformed with pREP41 (φ) or pREP41+Atf1. For overexpression, cells were grown in EMM-Leucine for 24 h at 30°C and then processed for live imaging. Bar represents 10μm. (B) Fixed cells were processed for microscopy after staining separately with DAPI (2µg/ml) and Calcofluor (2µg/ml) separately. Bar represents 10µm. (C) For overexpression, cells were grown in EMM-Leucine and growth was monitored for 5 days by measuring absorbance or optical density (OD) at 595 nm. The graph represents mean±s.e.m of 3 independent experiments. (D) *wt* and *Δpcr1* cells transformed with pREP41 (φ) or pREP41+Atf1 were grown in the presence of 20 µM Thiamine, and then washed to remove Thiamine and serial dilutions were spotted onto EMM-Leu plates. The plates were incubated at the indicated temperatures for 4 days before being photographed. All data are representative of 3 independent experiments.

### Changes in transcriptome and Spc1 activity are responsible for the observed morphogenetic transition

The bZIP domain of Atf1 and Pcr1 mediates their dimerization after which they can interact with target promoters either as heterodimers or homodimers. Dimerisation with Pcr1 is known to regulate the promoter specificity of Atf1 [26]. It is therefore logical to expect that the global transcription profile of the elongated cells that we found would be significantly different from the control cells and that such differences may be associated with the observed morphogenetic transition. To confirm such possibilities we repeated our experiments by overexpressing a truncated version of Atf1, namely Atf1ΔbZIP containing the N-terminal 1-474 residues and lacking the C-terminal bZIP domain. We found that this mutant when overexpressed in *Δpcr1* cells, failed to trigger the morphogenetic transition observed when full length Atf1 was overexpressed (Figure 2A). RNA-Seq analysis (data available at NCBI vide accession number GSE284036) of the *Δpcr1* cells where Atf1 had been overexpressed identified 105 genes whose expression was upregulated (cutoff ≥ 1.5) due to Atf1 overexpression (compared with *Δpcr1* cells transformed with empty vector) while only 3 genes were found to be downregulated (cutoff ≤ 0.75). These results are represented in Supplementary Table 1. 32 of these genes have been earlier reported to be upregulated in *wt* cells upon Atf1 overexpression and 9 of these genes were earlier reported to be downregulated in *wt* cells upon Atf1 overexpression. [It may be noted here though that the earlier reports are from cells in *h^-^* background while these current observations are from *S. pombe* cells in *h^90^* background]. Functional clustering analysis of these 105 genes using DAVID [40, 41] revealed that many of them are associated with important metabolic pathways in *S. pombe* cells (Figure 2B). These included genes involved in carbon metabolism, signalling, redox homeostasis, cell adhesion, flocculation, iron metabolism, etc (See Supplementary Table 1).

**Figure 2.**
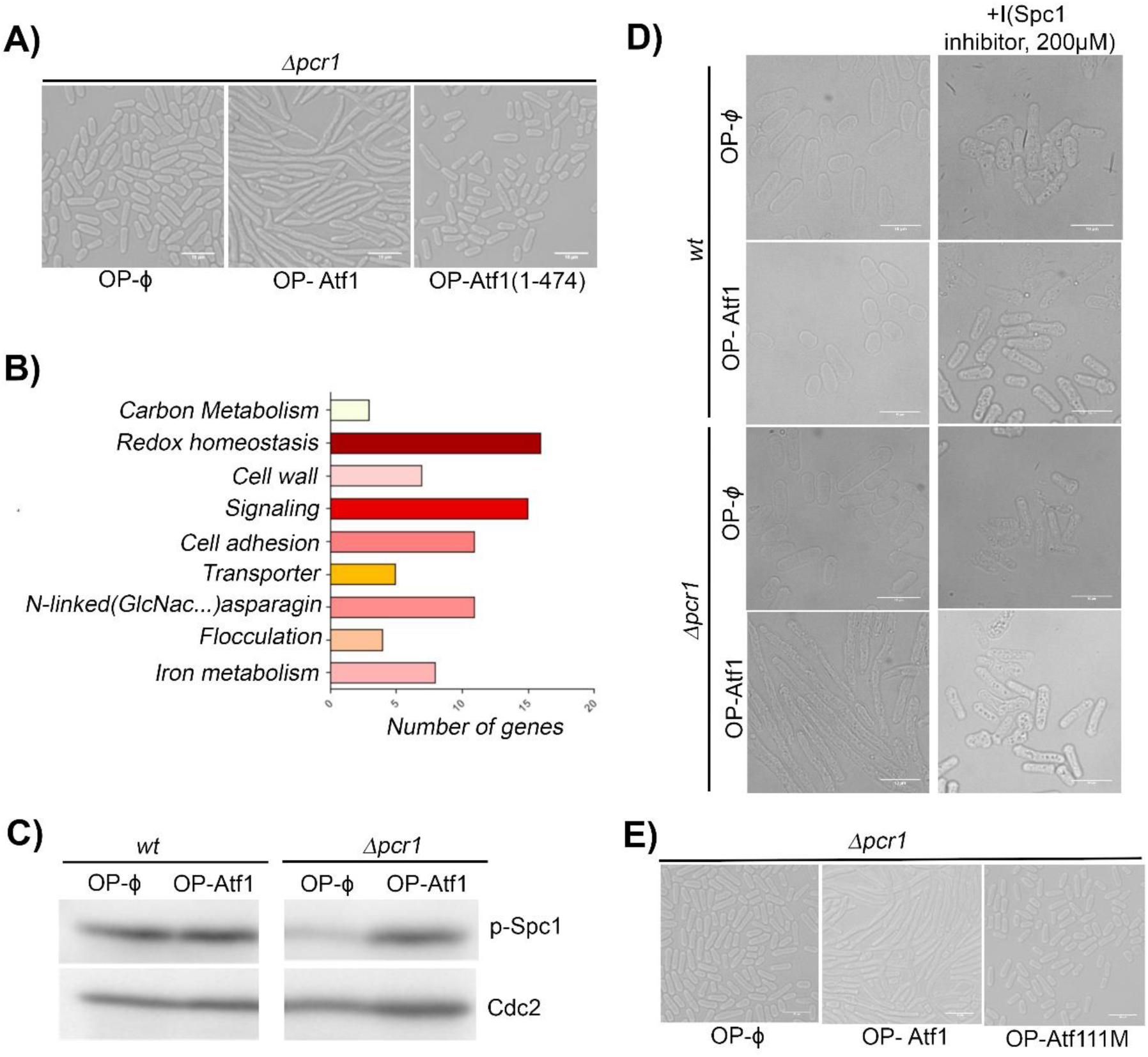
Changes in transcriptome and Spc1 activity are responsible for the observed morphogenetic transition. (A) Bright-field images of *Δpcr1* cells transformed with pREP41 (φ) or pREP41+Atf1 and pREP41+Atf1-11M. For overexpression, cells were grown in EMM-Leucine for 24 h at 30°C and then processed for live cell imaging. Bar represents 10 μm. (B) DAVID Annotation Tool [40,41] was used for functional clustering of genes upregulated upon Atf1-OP in *Δpcr1* background. Significant clusters include carbon metabolism, iron metabolism, redox homeostasis, etc. (C) Spc1 activity was observed upon Atf1-overexpression in *wt* and *Δpcr1* cells by checking the p-Spc1 levels. Cdc2 levels were used as loading control. (D) Bright-field images of *wt* and *Δpcr1* cells transformed with pREP41 (φ) or pREP41+Atf1 in presence and absence of Spc1 inhibitor (SP600125). For overexpression, cells were grown in EMM-Leucine for 24 h at 30°C with and without SP600125 and then processed for live cell imaging. Bar represents 10 μm. (E) Bright-field images of *Δpcr1* cells transformed with pREP41 (φ) or pREP41+Atf1 and pREP41+Atf1-11M. For overexpression, cells were grown in EMM-Leucine for 24 h at 30°C and then processed for live cell imaging. Bar represents 10 μm. All data are representative of 3 independent experiments.

Interestingly, yeast to hyphal transition is a property of many fungal pathogens and iron metabolism and redox homeostasis have significant roles in maintaining the pathogenicity and infectivity of fungal pathogens [42, 43, 44, 45]. The dimorphic phenotype of *S. pombe* cells that we observed in our experimental conditions and also the fact that they show variations in expression of genes, involved in processes, which have earlier been implicated in pathogenesis mechanisms in other fungi, reveal the possibility of a possible association between these metabolic processes and the morphology of fungal cells.

Analysis of the transcriptome revealed that 39% of the genes upregulated upon Atf1 over-expression in *Δpcr1* cells [See Supplementary Table 1] were previously known targets of Spc1 i.e. requiring Spc1 activity for regulation of their expression [46]. Regulation of Atf1 by the stress responsive Mitogen Activated Protein Kinase (MAPK) Spc1 is known to be important for its transcription related functions [25, 47, 48, 49, 50]. So from the RNA-Seq data, it seemed that Spc1 activity might be important for the observed morphogenesis in our experiments. Indeed we found that overexpression of Atf1 in *Δpcr1* cells resulted in increased Spc1 phosphorylation (Figure 2C). Since phosphorylated form of Spc1 is its active form, this observation indicates that Spc1 activity is high in the cells exhibiting the elongated morphology. We repeated our Atf1 over-expression experiments in the presence of a Spc1 inhibitor, namely SP600125. We found that treatment with SP600125 completely abolished the morphogenetic change associated with Atf1 over-expression in *Δpcr1* cells (Figure 2D). These results indicate that Spc1 activity is important for the Atf1 dependent morphogenetic transition of *S. pombe* cells. We then wanted to know whether phosphorylation of Atf1 by Spc1 is actually an important part of the mechanism triggering the morphogenesis. To investigate this, we repeated our experiments with a mutated non-phosphorylatable version of Atf1, namely Atf1-11M [47] in which all the predicted MAPK target sites are mutated. We found that over-expression of the non-phosphorylatable Atf1-11M mutant in *Δpcr1* cells, failed to trigger the morphogenetic transition that we were studying (Figure 2E).

Taken together these results indicate that the changes in global gene expression profile, increased Spc1 activity and phosphorylation of Atf1 possibly by Spc1 are all important for the mechanism of the observed morphogenesis of *Δpcr1* cells overexpressing Atf1.

### Alteration of iron homeostasis leads to significant change in morphogenesis of these cells through modulation of MAPK activity

An association between iron homeostasis and yeast to hypha transition has been reported earlier in pathogenic fungi [51]. So the presence of a cluster of genes associated with iron metabolism amongst the differentially expressed genes in *Δpcr1* cells overexpressing Atf1 intrigued us. We repeated our experiments separately in media containing either low (0.05μM.) or high (1.5μM) iron concentration. To our surprise we found that in both these conditions the extent of morphogenesis was less than that observed for cells growing in media containing optimal (0.74μM) iron concentration (Figure 3A). Specifically, we found that, percentage of elongated cells appears significantly to be less when *Δpcr1* cells overexpressing Atf1 are grown in low iron conditions and the phenotype seems somewhat heterogeneous. However when these cells were grown in media containing high iron conditions, the cells appeared a little less elongated than the ones growing in media with optimal iron concentration i.e. the phenotype appeared to be intermediate between normal cell length and filamentous cell length. The hits of the RNA-Seq included iron homeostasis associated genes like, like *fio1^+^*, *frp1^+^*, *fip1^+^* that are all important for growth in low iron conditions [52, 53, 54, 55].

**Figure 3.**
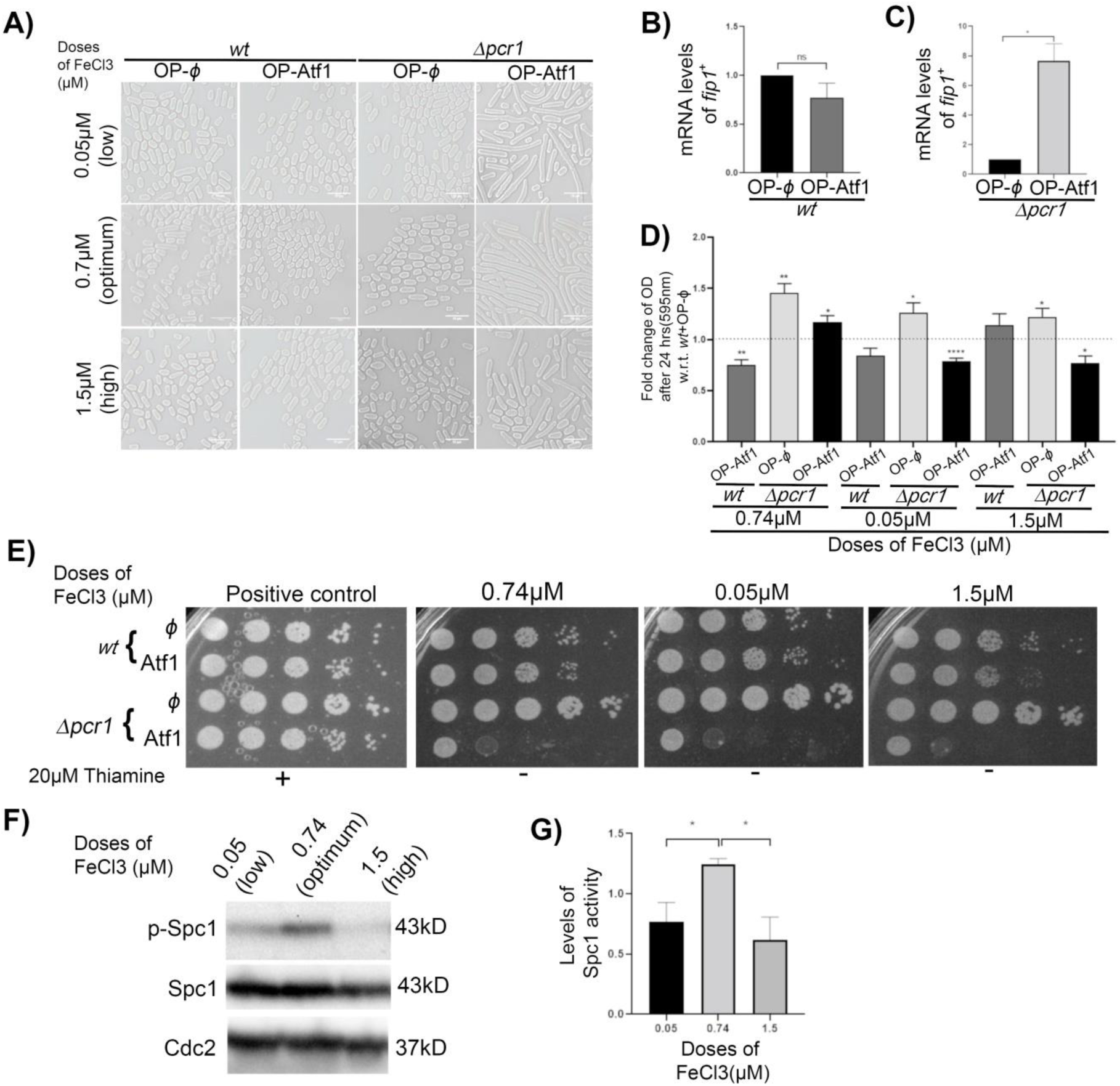
**Alteration of iron homeostasis leads to significant change in morphogenesis of these cells through modulation of MAPK activity**. (A) Bright-field images of *wt and Δpcr1* cells transformed with pREP41 (φ) or pREP41+Atf1 in low i.e 0.05μM FeCl_3,_ optimal i.e. 0.74μM FeCl_3_ and high i.e. 1.5μM FeCl_3_. For overexpression, cells were grown in EMM-Leucine containing the indicated doses of FeCl_3_ for 24 h at 30°C and then processed for live imaging. Bar represents10 μm. (B and C) Quantitative RT-PCR analysis of expression of genes involved in iron uptake in *wt* and *Δpcr1* cells transformed with pREP41 (φ) or pREP41+Atf1. 18S rRNA levels were used for normalization. Bars represent mean±s.e.m of three independent experiments. Statistical analysis was done using Graph Pad Prism application. t-test was used to evaluate the significance of data. *, P ≤ 0.05. (D) The growth of cells in optimum, low and high iron concentration (0.74μM, 0.05uM and 1.5uM FeCl_3_) after 24hrs of over-expression by measuring absorbance or optical density (OD) at 595nm and normalizing with the same measured before overexpression (0 hr). The fold change in growth increase w.r.t that observed for *wt*+OP-ϕ cells under similar conditions was calculated and plotted. Bars represent mean±s.e.m of three independent experiments. Statistical analysis was done using Graph Pad Prism application. Paired t-test was used to evaluate the significance of data. *, P ≤ 0.05; **, P≤ 0.01; ****, P≤ 0.0001 (E) *wt* and *Δpcr1* cells transformed with pREP41 (φ) or pREP41+Atf1 were grown in the presence of 20 µM Thiamine, and then washed to remove Thiamine and serial dilutions were spotted onto EMM-Leucine plates. The plates were incubated at the indicated temperatures for 4 days before being photographed. (F) Spc1 activity was observed upon Atf1-overexpression in *Δpcr1* cells growing in media containing the indicated doses of FeCl_3_ by checking the p-Spc1 levels. Cdc2 levels were used as loading control for normalisation. (G) Band intensities were quantified using ImageJ [68] and fold change of p-Spc1/total Spc1 ratios were determined. Bars represent mean±s.e.m of 3 independent experiments. Statistical analysis was done using Graph Pad Prism application. Unpaired t-test was done. *, P≤0.05. All data are representative of 3 independent experiments.

Fip1 is a transmembrane iron transporter which is also involved in reductive iron assimilation [56] via binding with Fio1 which is a plasma membrane iron transport multicopper oxidase.

Its expression is known to be induced during iron starvation [56], and it helps in increasing iron uptake during low iron conditions.

We validated the RNA-Seq results with qPCR and found that Atf1 overexpression does not affect *fip1^+^* expression in *wt* (Figure 3B) cells but it can significantly upregulate *fip1^+^* expression in *Δpcr1* cells (Figure 3C). These results indicate that presence of Pcr1 restricts the ability of Atf1 to activate *fip1^+^* transcription. Taken together with the earlier known roles of *fip1^+^*, this indicates towards a possible higher accumulation of iron in *Δpcr1* cells overexpressing Atf1. These results also indicate that the cellular iron levels are associated with the observed morphogenesis. The mechanistic connection between iron accumulation and morphogenesis however remains unclear.

We also looked at the growth of the cells in different iron conditions. The fold change in OD of the cultures after 24 hours of incubation in EMM-Leu media containing the indicated doses of iron in absence of Thiamine was calculated and normalized using the fold change seen for the *wt* cells transformed with empty vector and grown under similar conditions (Figure 3D). An interesting pattern that emerged was that in all the conditions deletion of *pcr1^+^*led to increased cell proliferation. Overexpression of Atf1 was however found to oppose this increase in normal to high iron conditions and to a lesser extent in low iron conditions. It is possible that since *Δpcr1* cells overexpressing Atf1 have high *fip1^+^* levels they might be accumulating iron more and in media containing normal to high iron concentrations, this increased accumulation of iron could be slightly toxic and thus contributing to decreased proliferation. Interestingly in both low and high iron conditions the effect of Atf1 overexpression on cell proliferation was completely opposite in *wt* and *Δpcr1* cells. In optimal iron conditions however overexpression of Atf1 affected *wt* and *Δpcr1* cells similarly. However, when we compared the growth of all these cells in media containing different iron concentration we did not find any significant differences (Figure 3E).

As we had found evidence that Spc1 activity was important for the morphogenesis, we then checked the same in *Δpcr1* cells overexpressing Atf1 when they were grown in media containing different iron concentration (Figure 3F, 3G). Interestingly, we once again found that the cells which were grown in optimal iron conditions and showed the maximum extent of morphogenesis, also had the highest levels of phosphorylated Spc1 (Figure 3F, 3G). Once again therefore, the results indicate a strong association between Spc1 activity and the morphogenesis of *Δpcr1* cells overexpressing Atf1.

### The morphogenesis of *Δpcr1 S. pombe* cells overexpressing Atf1 is also dependent on availability of glucose

Availability of glucose is a very important requirement for proper cell cycle progression in *S. pombe* and the cell cycle progression rates vary depending upon glucose concentration in the growth media [57, 58]. As earlier cited the cells get arrested when shifted to a respiratory media, glycerol for few hours but then they overcome the delay at steady rate [59]. *S. pombe* cells have known to grow in other carbon sources like maltose, fructose and galactose but little is known about their metabolism [60]. Since one of the gene clusters identified from our RNA-sequencing data was related to carbohydrate metabolism, we next investigated the influence of carbon sources on the observed morphogenesis of *Δpcr1 S. pombe* cells overexpressing Atf1. We observed significant differences in the morphogenesis of the cells when shifted to different carbon sources like 2% maltose, 2% fructose, 2% galactose and 2% glycerol-1% ethanol media (Figure 4A). Cells growing in media containing maltose and fructose showed an intermediate phenotype with slightly shorter filamentous forms compared to those growing in media containing 2% glucose. This reduction in the extent of morphogenesis was more pronounced in presence of fructose than in maltose. It may be noted here that maltose can be hydrolyzed to yield glucose and ultimately the cells can follow a similar metabolic pathway for energy production. Metabolism of fructose however requires a different metabolic pathway [61]. Interestingly shifting the cells to respiratory media containing either galactose or glycerol-ethanol as the carbon source completely abolished the morphogenetic transition. These results unravel the interesting fact that the metabolic pathways for energy production adopted by the cell can completely dictate the cell fate in terms of the observed morphogenesis. *S. pombe* cells growing in fructose are also known to have a tendency to shift their metabolic activity to a slightly respiratory mode [61] and that might contribute to the observed intermediate phenotype of those cells.

**Figure 4.**
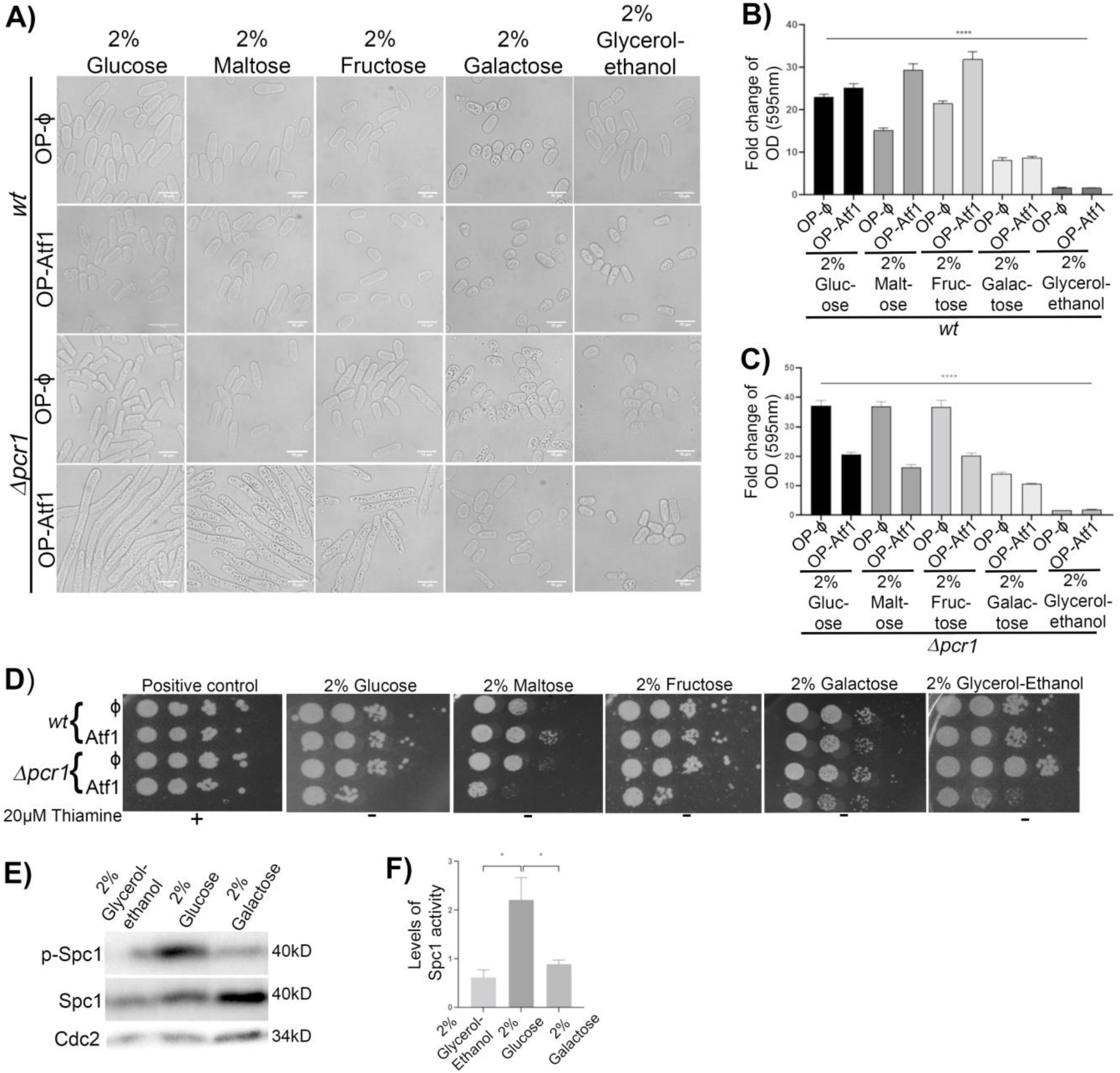
The morphogenesis of *Δpcr1 S. pombe* cells overexpressing Atf1 is also dependent on availability of glucose. (A) Bright-field images of *wt and Δpcr1* cells transformed with pREP41 (φ) or pREP41+Atf1 and for overexpression cells were grown in EMM-Leucine in 2% glucose, 2% maltose, 2% fructose, 2% galactose and in 2% glycerol-ethanol media for 24 h at 30°C and then processed for live imaging. Bar represents 10 μm. (B) The extent of growth of cells, *wt* +OP-ϕ and *wt* +OP-Atf1 in liquid media after 24h of over-expression in 2% glucose, 2% maltose, 2% fructose, 2% galactose and 2% glycerol-ethanol media were measured at 595nm, normalized by the OD at 0hr. Bar represents mean± s.e.m of 3 independent experiments. A one-way Anova was done to compare the ten groups and revealed a significance difference between their means[****, P≤ 0.0001]. Post hoc analysis using Tukey’s multiple comparison tests was done (See Supplementary Table 2) (C) The extent of growth of cells, *Δpcr1* +OP-ϕ and *Δpcr1* +OP-Atf1 in liquid media after 24h of over-expression in 2% glucose, 2% maltose, 2% fructose, 2% galactose and 2% glycerol-ethanol media was measured at 595nm, normalized by the OD at 0hr. Bar represents mean± s.e.m of 3 independent experiments. A one-way Anova was done to compare the ten groups and revealed a significance difference between their means[****, P≤ 0.0001]. Post hoc analysis using Tukey’s multiple comparison tests was done (See Supplementary Table 2) (D) *wt and Δpcr1* cells transformed with pREP41 (φ) or pREP41+Atf1 were grown in the presence of Thiamine, and then washed to remove Thiamine and serial dilutions were spotted onto EMM-Leu plates. The plates were incubated at the indicated temperatures for 4 days before being photographed. (E) Spc1 activity was observed upon Atf1-overexpression in *Δpcr1* cells growing in media containing the indicated carbon sources by checking the p-Spc1 levels. Cdc2 levels were used as loading control for normalisation. (F) Band intensities were quantified using ImageJ [68] and fold change of p-Spc1/total Spc1 ratios were determined. Bars represent mean±s.e.m of 3 independent experiments. Statistical analysis was done using Graph Pad Prism application. Unpaired t-test was done. *, P≤0.05. All data are representative of 3 independent experiments.

We also compared the growth of these cells. We found that overexpression of Atf1 rescues the growth defect of *wt* cells in media containing 2% maltose and 2% fructose (Figure 4B). In respiratory media however no such rescue was seen in *wt* cells. *Δpcr1* cells were found to grow equally well in media containing glucose, fructose or maltose and overexpression of Atf1 was surprisingly found to decrease the growth of these cells (Figure 4C). Similar trend was also seen in the differences in viability of these cells (Figure 4D). Interestingly, *Δpcr1* cells overexpressing Atf1 grown in respiratory media, where the morphogenesis was found to be abolished, were found to be much more viable than the filamentous counterparts growing in fermentative media (Figure 4D). These results reveal that the morphogenetic transition is directly associated with decreased viability.

We also observed that the levels of phosphorylated Spc1 were lower when cells were grown in galactose or glycerol-ethanol containing media (Figure 4E, 4F). So the MAPK Spc1 definitely plays a crucial role in regulation of the morphogenesis of *Δpcr1* cells overexpressing Atf1.

## Discussion

Yeast to hyphal transition has been seen earlier in many pathogenic fungi. In case of non-pathogenic fission yeast cells, *S. japonicus* has also been observed to be dimorphic with pseudohypha formation, high growth rate, and aberrant septation [62]. Here we report that overexpression of the bZIP transcription factor Atf1 ( human ATF2 homolog) in absence of its binding partner and functional regulator Pcr1 ( human CREB homolog) results in a transition from yeast to filamentous form with cells becoming hyper elongated with reduced septation and aberrant nuclear morphology. We also show that the molecular mechanism associated with this morphogenesis involves Spc1 activation and the phosphorylation of Atf1 and global gene expression changes associated with the transcription regulating functions of Atf1. Inhibition of either Spc1 activity or Atf1 phosphorylation or the DNA binding of Atf1, all these result in abolishment of the morphogenetic transition. We also show that the morphogenesis is associated with optimal iron concentration in the growth media and also that shifting the cells to respiratory media abolishes this morphogenetic transition of *Δpcr1* cells overexpressing Atf1. More importantly we also show that loss of this transition is associated with better survival of these cells. Growth conditions leading to dimorphisms are well known in yeast cells. This is the first time however that we show that changes in levels of bZIP transcription factors can also cause the same especially in a non-pathogenic yeast.

Tissue differentiation in higher eukaryotes is known to be associated with morphogenesis [63]. Our results show a relationship between energy metabolism and morphogenesis in *S. pombe* cells thus indicating that the complex processes involved with differentiation and development in multicellular organisms have their origins in simple unicellular eukaryotes also. bZIP transcription factors are known to have a strong influence on morphogenesis and development in plants [64]. ATF2, the human homolog of *S. pombe* Atf1 has been shown to be important for several developmental processes and diseases in mammals [65, 66]. Our findings on the association of alteration of bZIP transcription factor levels with morphogenesis and cell fate establish the fundamental nature of such mechanisms and can in future contribute towards development of a genetically amenable model for studying many complex processes associated with these phenomena. Moreover, these results can also be used to understand mechanisms associated with fungal pathogenesis and thus this non-pathogenic model can be useful in that regard too. Morphogenetic transitions are intricately related to evolution and further investigation of the mechanism and processes reported in this paper in other fungal species and well as higher eukaryotic systems can lead to a deeper understanding of the relationship between morphogenesis and evolution.

## MATERIALS AND METHODS

### Strains, media and growth conditions

The strains used in this study are listed in Table 1. In general cells were grown overnight in Edinburgh Minimal Medium (EMM) without Leucine, containing 2% Glucose as carbon source and supplemented with 20 μM Thiamine. Subsequently, they were washed and resuspended at an OD of 0.05 into media containing no Thiamine to allow overexpression of the indicated gene and grown for another 24 hrs or as long as indicated otherwise at 30ᵒC. For experiments involving variations in iron concentration, the mineral stocks (10000X) for EMM were prepared with 0.5mM and 15mM FeCl_3_ instead of the usual 7.4mM while other components of the stock remained the same. EMM media was supplemented with three different types of mineral stocks at a final concentration of 1X. For experiments involving different carbon sources, the EMM was supplemented with either of 2% Maltose, 2% Fructose, 2% Galactose, 2% Glycerol-1% Ethanol instead of 2% glucose.

**Table 1.** List of strains/plasmids in the study.

| **Strain and plasmid no.** | **Genotype** | **Source** |
| --- | --- | --- |
| **GSY500** | *h<sup>90</sup> leu1-32ura4D-18* | Nakamura-Shimoda |
| **GSY531** | *h<sup>90</sup> ade6 leu1ura4 pcr1::ura<sup>4+</sup>* | Elena Hidalgo |
| **pGS017** | pREP41 | Yeast Genetic Resource Centre |
| **pGS018** | pREP41+Atf1 | Labstock |
| **pGS045** | pREP41+Atf1ΔbZIP | Labstock |
| **pGS139** | pREP41+Atf1-11M | Labstock |

### Schizosaccharomyces pombe transformations

1ml of an overnight *S. pombe* culture in YES (Yeast Extract with supplements) was harvested and then resuspended in 0.5 ml PEGLET (10 mM Tris [pH 8], 1 mM EDTA [pH 8], 0.1 M lithium acetate, 40% polyethylene glycol [PEG]). 5µg of denatured salmon sperm DNA (10 mg/ml) was added to it. 1µg of the purified plasmid DNA was then added to this mixture and allowed to stand overnight at room temperature, after which the cells were resuspended in 150 μl YES and spread onto appropriate selection plates.

### Microscopy

70% ethanol was used for fixing the cells and imaging was done using the Olympus BX51 Fluorescence Microscope at 100X magnification for visualization of septa using Calcofluor (2µg/ml) staining and imaging of nuclei using DAPI (2µg/ml) staining. Live cell imaging was done using Olympus BX53 Microscope using the brightfield filter at 40X magnification using unstained cells. Live cell imaging for Figure 2D and 4A was done using Nikon Eclipse TX 2R at 100X magnification.

### Protein extraction and immunoblotting

Whole cell extracts were prepared as described before [67]. Cells were harvested and resuspended in 20% Trichloroacetic acid (TCA) followed by addition of glass beads and vortexed at maximum speed for five 1 min pulses. The lysate was then transferred to a microcentrifuge tube and then centrifuged at 13,000 rpm for 15 mins. The supernatant was discarded and the pellets were washed with 70% ethanol thrice. All steps were performed at 4°C and samples were kept on ice. Finally, the pellets were resuspended in Laemmli buffer and boiled for 5 min at 95°C. The samples were then loaded onto SDS–polyacrylamide gels. After transferring onto PVDF membrane, immunoblotting was performed using anti-Cdc2 (Sc-53217) antibody, anti-Phospho p38 (Cell Signalling Technology# 9211S, for Phospho-Spc1) and anti-Spc1 (labstock) antibodies at 1:1000 dilutions. Immunoblots were developed using Clarity™ Western ECL substrate (Cat No# 170-5060).

### Viability assays

Cells were grown in 2% glucose media (EMM with Thiamine), followed by washes and then resuspended to make the final concentration at 1 OD. Tenfold serial dilution of cultures of exponentially growing cells was made. These dilutions (5 μl) were spotted onto suitable media plates. Photography of these plates was performed after incubation for 4 days at a particular temperature.

### cDNA preparation for quantitative RT PCR

Total cellular RNA was isolated as mentioned above. DNAseI (Thermo Scientific #EN0521) treatment was done to remove genomic DNA contamination. 1µg of the DNaseI treated RNA was used for cDNA preparation using random hexamers and M-MuLV reverse transcriptase (Thermo Scientific RevertAid First Strand cDNA Synthesis Kit, Cat# K1622). Quantitative PCR was performed using CFX96 Real Time System of BioRad using SYBR Green reagent (TB Green Premix Ex Taq Tli RNase H Plus, Cat# RR420A). Melt curve analysis was done to confirm the absence of primer dimers and non-specific amplification.18S rRNA levels were used for data normalisation. Sequence of primers (in 5’ to 3’ direction) used are as follows:

*fip1^+^* forward: TCTCATCGTTGGTTGTCTTATC

*fip1^+^* reverse: CGGAGATAAGGTAGAGGATACA

*18S rRNA^+^* forward: AAGACCGACCTTCCTTACTTTG

*18S rRNA^+^* reverse: GCCGGTCTCTGCTGTAATAAT

### Statistical analysis

Statistical analysis of all representative data was done using GraphPad Prism version 8.0.2 for Windows [68]. All bar graphs represent mean±s.e.m. For evaluation of statistical significance of all quantitative data, a one-tailed *t*-test was performed when comparing two groups of data and one-way ANOVA with post hoc analysis was done when comparing more than two groups of data. *P*-value was used to estimate the significance of the results: \*\*\*\**P*<0.0001; \*\*\**P*<0.001; \*\**P*<0.01; \**P*<0.05; ns, not significant.

## Author contributions

G.S. and S.D. designed the experiments. S.D, S.D and J.B. have performed the experiments. S.D and G.S analysed the data. S.D and G.S contributed in writing of the manuscript.

## Abbreviations

MAPK: Mitogen Activated Protein Kinase
cAMP: cyclic AMP
NGS: Next Generation Sequencing
DAVID: Database for Annotation, Visualization and Integrated Discovery
RT-PCR: Real-time-polymerase chain reaction
EDTA: Ethylenediamine tetraacetic acid
PVDF: Polyvinylidene fluoride
ECL: Electrochemiluminescnence

## Acknowledgements

Transcriptome sequencing and analysis service has been obtained from Agrigenome Labs Private Limited, Cochin. The authors thank DBT-IPLS-CU, UGC-CAS, Central Instrument Facility, University of Calcutta, DST-FIST programme of Department of Biochemistry, University of Calcutta for infrastructural support. S.D. thanks WB-DBT [Ref No. 56(Sanc.)-BT/(Estt.)/RD-17/2017 dated 13/08/2018] for Fellowship. The authors acknowledge Science and Technology and Biotechnology Department, Govt. of WB [2403(Sanc.)/STBT-13015/8/2020-BT-SEC dated 20/03/2024] for funding. We are grateful to Prof. Anindita Seal, Department of Biotechnology, University of Calcutta for allowing us to use the Brightfield microscope.

## Notes

### Competing Interest Statement

The authors have declared no competing interest.

